# Voluntary alcohol intake alters the motivation to seek intravenous oxycodone and neuronal activation during the reinstatement of oxycodone and sucrose seeking

**DOI:** 10.1101/2023.07.20.549769

**Authors:** Courtney S. Wilkinson, Harrison L. Blount, Shane Davis, Giselle Rojas, Lizhen Wu, Niall P. Murphy, Marek Schwendt, Lori A. Knackstedt

**Author notes:** **Corresponding author:** Lori A. Knackstedt University of Florida 114 Psychology 945 Center Dr. Gainesville, FL 32611.

## Abstract

Opioid-alcohol polysubstance use is prevalent and worsens treatment outcomes. Here we assessed whether co-consumption of oxycodone and alcohol would influence intake of one another, demand for oxycodone, and the neurocircuitry underlying cue-primed reinstatement of oxycodone-seeking. Male and female rats underwent oxycodone intravenous self-administration (IVSA) with access to either alcohol (20% v/v) and water or only water immediately after the IVSA session. Next, economic demand for intravenous oxycodone was assessed while access to alcohol and/or water continued. Control rats self-administered sucrose followed by access to alcohol and/or water. Rats underwent extinction training and brains were processed for c-fos mRNA expression immediately following a cue-primed reinstatement test. While both sexes decreased oxycodone intake if they had access to alcohol, and decreased alcohol intake if they had access to oxycodone, female oxycodone+alcohol rats exhibited decreased demand elasticity for intravenous oxycodone and increased cue-primed reinstatement while male rats did not. Spontaneous withdrawal signs were correlated with oxycodone intake while alcohol intake was correlated with anxiety-like behavior. Alcohol consumption increased the number of basolateral and central amygdala neurons activated during sucrose and oxycodone reinstatement and the number of ventral and dorsal striatum neurons engaged by sucrose reinstatement. Nucleus accumbens shell dopamine 1 receptor containing neurons displayed activation patterns consistent with oxycodone reinstatement. Thus, alcohol alters the motivation to seek oxycodone in a sex-dependent manner and alters the neural circuitry engaged by cue-primed reinstatement of sucrose and oxycodone-seeking.

## Introduction

From 2002 to 2012 there was a 15-fold increase in the prevalence of persons with alcohol use disorder (AUD) also reporting opioid use disorder (OUD), from 2.7% to 42% [1][2]. Co-occurring alcohol use hastens the time from onset of opioid use to OUD diagnosis [3] and predicts poor treatment outcomes [4]. Alcohol is found with opioids in the blood of 14-16% of opioid overdose patients [5–7]. Oral oxycodone administered prior to oral alcohol increases ratings of drug liking and willingness to take these drugs again relative to placebo; with no effects of either drug alone on these ratings [8]. Thus, opioid-alcohol polysubstance use (PSU) is prevalent and clinically relevant, as it increases the probability of future use and worsens OUD trajectories.

Animal models of opioid-alcohol PSU are essential for the identification of neuroadaptations unique to PSU. Intravenous self-administration (IVSA) is the most widely used animal model of drug-seeking. Of the 1800+ publications on opioid IVSA (PubMed), none involve the consumption of alcohol. To begin to investigate opioid-alcohol PSU, here we assess the ability of sequential self-administration of oxycodone and alcohol to influence the motivation to seek and consume both drugs in rats. We chose oxycodone, as opposed to other opioids, because of its prevalent use and greater association with overdose deaths relative to other opioids like heroin [9,10]. There is currently little data regarding the patterns of opioid-alcohol PSU employed (i.e. sequential, simultaneous), with the exception of reports that 16-20% of opioid users consume alcohol on the same day [11,12], in agreement with the prevalence of alcohol detected in the blood of opioid overdose victims. Thus, we designed a sequential model of PSU in which both drugs are consumed on the same day with overlapping effects. Alcohol access was provided after operant sessions to allow for testing of the motivation to seek/take oxycodone in the undrugged condition.

Opioid withdrawal is associated with increased motivation to seek opioids [13]. In rodents, somatic signs of withdrawal (SSW) are consistently observed following non-contingent administration and voluntary self-administration of opioids, both when “precipitated” with an opioid antagonist and spontaneously [14–17]. Non-contingent alcohol exposure reliably produces SSW in withdrawal, as does voluntary alcohol intake in mice [18]. In addition to producing SSW, early abstinence (24-36 hrs) from voluntary alcohol consumption results in anxiety-like behavior in the elevated plus maze and other assays [19–21]. Interestingly, oxycodone is anxiolytic in male and female rats [22]. Additive or synergistic pharmacological effects of opioids and alcohol may underlie the motivation for co-consumption.

We hypothesized that the PSU condition would result in drug intake that would have additive effects on SSW and anxiety-like behavior. We also hypothesized that different patterns of neuronal activity would be observed following reinstatement of oxycodone in the PSU and oxycodone-only conditions, as was found following cocaine-only and sequential cocaine+alcohol IVSA [23]. Specifically, we predicted that increased oxycodone-seeking in the PSU condition would be accompanied by increased c-fos expression in cell populations established to be involved in reinstated-drug seeking: neurons that release glutamate into the striatum, namely prelimbic (PL) cortex and basolateral amygdala (BLA) neurons expressing the vesicular glutamate transporter 1 (vGlut1), and the downstream dopamine 1 receptor (D1)-expressing cells of the nucleus accumbens (NA) core and the dorsolateral striatum (dSTR) [24–29]. We hypothesized that increased reinstatement would be accompanied by decreased c-fos expression in circuits that inhibit drug-seeking, namely glutamate neurons of the infralimbic (IL) cortex that project to the NA shell [30]. To test these hypotheses, we employed a 3-phase model in which rats first established oxycodone IVSA with low-effort required to obtain drug, followed by determination of the elasticity of demand for IV oxycodone upon increases in the response requirement necessary to obtain a single infusion. During both phases, rats had access to alcohol and/or water in the home cage immediately following the operant sessions. The last phase was the extinction and cue-primed reinstatement of oxycodone-seeking followed by c-fos expression analysis. We used classic inferential statistics to test the hypothesis that alcohol co-use would alter oxycodone intake and seeking, as well as anxiety-like behavior, withdrawal and reinstatement-induced c-fos expression. We also used machine-learning clustering technique to examine the ability of sex and alcohol intake to alter the relationship between these dependent variables.

## Materials and Methods

### Subjects

Sprague-Dawley rats (n=60; half male; 8 weeks old; Charles River, Raleigh, NC) were single housed on a reversed 12-hr light cycle (lights on at 7 am) in a temperature-controlled vivarium. Procedures were approved by the University of Florida IACUC and are reported according to ARRIVE guidelines.

### Drugs

Oxycodone HCl (Sigma Aldrich) was prepared in 0.9% physiological saline at the concentration of 0.4 mg/mL for male rats and 0.32 mg/mL for females. Alcohol (100%; Fisher Scientific) was diluted with tap water to 20% (v/v).

### Surgery

Rats were anesthetized using ketamine (87.5 mg/kg, IP) and xylazine (5 mg/kg, IP). Ketorolac (2 mg/kg, IP) was administered post-operatively and 3 days following surgery for analgesia. Catheters (SILASTIC silicon tubing, ID 0.51 mm, OD 0.94 mm, Dow Corning, Midland, MI) were implanted in the jugular vein, secured with sutures, and passed subcutaneously between the shoulder blades to exit through the skin on the back. Catheter tubing was connected to a stainless-steel cannula (Plastics One, Roanoke, VA, USA) embedded in a rubber harness (Instech, Plymouth Meeting, PA, USA) that was worn for the duration of self-administration. The antibiotic cefazolin (100 mg/kg) was administered IV (0.1 mL) for 3 days post-surgery. Catheters were flushed with heparin (100 IU/mL; 0.1 mL) before and after each self-administration. Catheter patency was tested periodically with methohexital sodium (10 mg/mL; Eli Lilly, Indianapolis, IN, USA), which results in a temporary loss of muscle tone.

### Self-administration, demand analyses, extinction and reinstatement

One week after arrival, rats that would later have access to alcohol (ALC) following operant sessions were exposed to intermittent access to alcohol (IAA). Rats were provided 2-bottle choice for unsweetened alcohol in the home cage in five 24-hour sessions alternating with 24 hr periods without alcohol access (see timeline in Fig. 1a). Rats were then implanted with jugular catheters and permitted 6 days to recover. Thirty-six rats underwent oxycodone IVSA (0.1 mg/kg/infusion [31]) for 3 hr/day for 6 days on an FR-1 schedule of reinforcement, followed by 6 days on an FR-3 schedule. Self-administration took place in 2-lever operant chambers (Med Associates, Inc). Presses on the active lever delivered IV oxycodone (OXY) and drug-associated cues (stimulus light and 2900 Hz tone). There was a 20-sec time out following reinforcer delivery, during which time presses were recorded but reinforced. Presses on the inactive lever had no programmed consequences. Twenty-four rats (half male) self-administered 45 mg sucrose (SUC) pellets (BioServ) instead of oxycodone in the same manner. Immediately following the SA session, rats were placed into the home cage for 6 hr access to 2-bottle choice or water only (n=18 OXY and n=12 SUC in each condition).

**Fig. 1.**
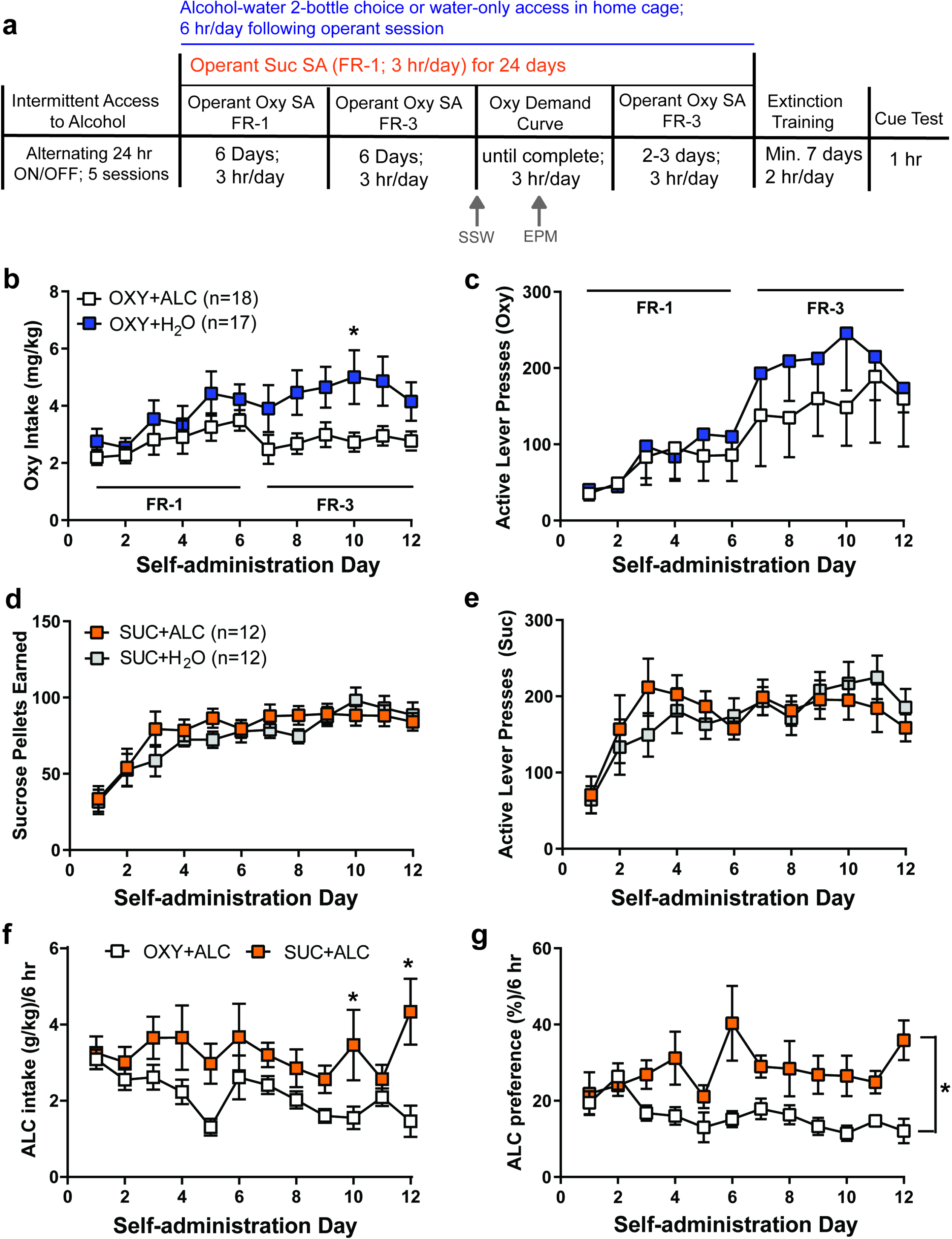
Alcohol and oxycodone influence the intake of one another. **a.** Timeline for the experiment. All rats experienced either 2-bottle choice or water access in the homecage. Rats self-administered either oxycodone (OXY) or sucrose (SUC) in the operant chamber for 12 days; OXY self-administering rats then underwent demand curve analyses, while the sucrose (SUC) self-administering rats did not. Alcohol/water access was provided daily during this time, immediately following the operant session. All rats underwent extinction training in the absence of alcohol access, followed by a cue-primed reinstatement test. There were no effects of sex on oxycodone, sucrose and alcohol self-administration and thus males and females are graphed together here. **b.** The OXY+H_2_O condition displayed greater oxycodone intake than the OXY+ALC condition on Day 10. **c.** Access to alcohol had no effect on active lever presses for oxycodone. **d**. Alcohol access did not influence sucrose intake or active lever presses (**e**) for sucrose. **f.** The SUC+ALC group consumed more alcohol than the OXY+ALC group on Days 10 and 12. **g**. There was an overall decrease in alcohol preference in the OXY+ALC group relative to the SUC+ALC group. * = p<0.05.

After 12 days of IVSA, demand for oxycodone was assessed. Demand curve sessions were identical to OXY IVSA training sessions, except that the FR requirement to attain an infusion of OXY was increased by quarter-and then eighth-log units (e.g. FR-3, FR-6, FR-10, FR-13) every second day. Access to alcohol and/or water continued as originally assigned. Sucrose rats did not undergo demand curve procedures, but self-administered sucrose on an FR-1 for 24 sessions. After completion of the demand procedure (earning 0 infusions on both days of a FR), rats re-established OXY IVSA on an FR-3 for 2-3 days until infusions were within 25% of the training average, followed by instrumental extinction training for a minimum of 7 days and until meeting criterion (<25% active lever presses during last day of IVSA). SUC rats underwent extinction training to the same criteria. Rats that met criteria by Day 16 were tested for cue primed reinstatement in a 1-hr test, during which the active lever yielded reinforcer-associated cues. Rats were killed immediately via rapid decapitation without anesthesia, brains flash frozen and stored at -80°C. This timing was chosen as it captures the peak of c-fos mRNA expression [32]. See Fig. 1a for timeline.

### Assessment of spontaneous withdrawal and anxiety-like behavior

After completing 12 days of self-administration, rats were assessed for spontaneous somatic signs of withdrawal (SSW) 20-22 hrs after the last self-administration session (Fig. 1a). Rats were placed into a clear plexiglass chamber (16” W x 16” D x 15” H) positioned under a camera for a 20-minute assessment. Videos were later rated by a researcher blind to condition. The following behaviors were quantified and summed for each rat: wet dog shakes, biting, grooming, jumping, and scratching. These behaviors are typically assessed for the assessment of withdrawal from opioids [17,33].

All rats completed 18 days of self-administration (FR-18). Anxiety-like behavior was assessed at 20-22 hrs withdrawal following the last FR-18 session. Rats were tested on the elevated plus maze (EPM) using an apparatus from Med Associates (St. Albans, VT, USA). The black plexiglass EPM had four arms (50 cm length x 10 cm width) raised 72 cm from the floor. Two open arms (1.5 cm high walls) and two closed arms (50 cm high walls) were joined by a center square platform (10 cm x 10 cm). The center platform was illuminated at 50 lux. Rats were individually placed on the center platform and given 5 min to explore the maze. Time spent in the open arms (OA), closed (CA), and number of OA and CA entries were measured using EthoVision XT 14 software (Noldus Information Technology, Leesburg, VA).

### Fluorescent in situ hybridization (FISH) and Imaging for the quantification of c-fos mRNA expression

Coronal sections (14 µM) were obtained on a Leica CM1950 cryostat and underwent FISH using probes for c-fos and D1 or vGlut1. Regions of interest included the PL and IL cortices, NA core (NAc) and shell (NAs), dSTR, BLA, and the central nucleus of the amygdala (CeA). Fluorescent in situ hybridization was performed using the RNAscope Multiplex Fluorescent Reagent Kit (Advanced Cell Diagnostics; Newark, CA, USA) according to the manufacturer’s instructions with some modifications (Shallcross et al., 2021). Slide-mounted frozen tissue was fixed for 15 minutes in 4% paraformaldehyde (PFA; 4°C, pH 7.45), and then dehydrated in an ethanol gradient (50%, 70%, 100%). Following protease digestion, slides were twice washed with PBS and hybridized with probes for c-fos in combination with either D1 (striatum) or vGluT1 (cortex and amygdala) for 2 hours at 40°C. The following probes (Advanced Cell Diagnostics) were used: c-fos (FOS, Cat #403591), D1 (Drd1a, Cat. # 317031), and vGlut1 (SLC17A7; Cat. #317001). In the PL, IL, and BLA c-fos and vGlut1 probes were used, as glutamate neurons in these regions projecting to the striatum have been shown to regulate the reinstatement of drug-seeking [34–36]. In the dSTR, NAc and NAs, c-fos and D1 probes were used, as D1- containing neurons have been shown to drive reinstated drug-seeking[37]. Antifadant mounting media containing DAPI counterstain was used.

### Image acquisition and quantification

A Leica DM6B microscope equipped with a sCMOS K5 camera and LASX software (v.3.7.423463) was used to acquire images using a 40x objective. In each region, 10 µm z-stacks were taken with 1 µm steps. Two sections were imaged from each brain region/rat. Colocalization and quantification of mRNA transcripts were done with CellProfiler (v. 4.0.7; http://cellprofiler.org/). mRNA expression was quantified within a 10-pixel region surrounding nuclei. The dependent measures were the number of c-fos+ cells and the number of c-fos/vGluT1 or c-fos/D1 doubled labeled cells.

### Statistical Analyses

Data were analyzed using SPSS (v.28, IBM) for 3-way ANOVAs and GraphPad Prism (v.9.4.1, GraphPad Software) for all other parametric tests and demand curve analyses. R (v. 4.2.2) was used to conduct the multi-dimensional scaling (MDS) analysis [38]. 3-way ANOVAs compared self-administration variables (Sex x Liquid x Time), alcohol intake/preference (Sex x Reinforcer x Time), and EPM and SSW dependent measures (Sex x Liquid x Reinforcer). Alcohol preference was computed by dividing the amount of alcohol consumed by the total amount of liquid consumed. Reinstatement of oxycodone-seeking was defined as a significant increase in active lever presses during the cue test compared to the last day of extinction; this comparison was made using paired-samples t-tests with the a priori hypothesis that all groups would reinstate. Significant interactions were followed by Sidak’s post-hocs.

Economic demand for oxycodone was assessed by plotting drug consumption (mean infusions averaged over the 2 days of each FR) as a function of FR and fitting the equation log*Q*=log(*Q*_0_)+k(e^-α*P*^ – 1)[39]. *Q* is the number of infusions at each FR requirement, or “price (*P*)”. *K* is a fixed scaling parameter representing the range of the dependent variable in logarithmic units, which for this data set was 4.031. *Q*_0_ is the y-intercept, and estimates drug consumption at an FR of 0 (i.e., when the drug is “free”). Alpha (*α*) is an index of elasticity of demand, which reflects the rate of change in consumption as a function of response requirement increases. Essential value, or α transformed, is calculated with the formula 1/(100αk^1.5^), and is the rate of change of the elasticity of demand [39]. P_max_ is calculated with the formula m/(Q_0_αk^1.5) where m = 0.084k + 0.65. Reinforcer intake at prices (FR values) lower than P_max_ are relatively stable, while intake at FR values higher than P_max_ diminish rapidly upon response requirement increases [39,40]. The extra sum-of-squares F-test was used as a nonparametric test of the null hypothesis that *α* did not differ between groups and 2-way (Liquid x Sex) ANOVAs compared *α*, *Q*_0,_ *α* transformed, and P_max_ between conditions [40].

mRNA expression was averaged between two images/region for each rat. For the OXY groups, a 3-way Sex x Liquid x Test ANOVA was conducted to identify regions which responded to the cue test. Expression in tested and untested rats was then analyzed separately with 2-way Sex x Liquid ANOVAs. Pearson’s correlations assessed the relationship between select dependent variables. Significant interactions were followed by Sidak’s post-hoc tests, correcting for multiple comparisons.

To examine the ability of Sex and Liquid to change the relationship between dependent variables, multidimensional scaling (MDS) analysis was used. MDS uses statistical approaches to graphically represent the similarity/dissimilarity between variables. Here, similarity was based on Pearsons correlations (computed via the R package *cor),* that were calculated within each condition (OXY+ALC male and female; OXY+H_2_O male and female; SUC+ALC and SUC+ H_2_O). Sucrose conditions were not examined by Sex due to the low n for many variables. Uncorrected p-values were calculated using *rcorr$p*. All correlation coefficients were converted to their absolute values (i.e., abs(r)), which was used to produce a distance matrix based on 1-abs(r) which was then subjected to MDS analysis. Subsequently, groups of correlated behaviors were identified using K-means clustering via the R function *<u>kmeans</u>* which applies a machine learning clustering algorithm to identify clusters of correlated variables. *Nbclust* was used to identify the optimal number of clusters and instructed to identify the optimum a range of 2-7 clusters. Variables included in this analysis included those that were found to be different by group/sex and in the case of variables being inherently related (e.g. time spent in the OA and time in CA; oxycodone intake and infusions), we selected only one representative variable. For sucrose rats, based on smaller n’s and a lack of an effect of sex on alcohol intake, reinstatement of sucrose-seeking, and c-fos expression, analyses were not separated by sex. Both two and three week totals for alcohol and oxycodone intake were used in the analysis because SSW were assessed after two weeks and EPM after three weeks. SSW could not be incorporated into the analysis for SUC rats, since the three rats without SSW ratings were some of those that were tested for reinstatement.

## Results

At the onset, OXY+ALC and OXY+H_2_O conditions comprised 18 rats, half of which were male. One male OXY+H_2_O died during IVSA and this data is excluded from analyses; 4 rats completed IVSA but died/lost patency during the demand curve (2 female OXY+ALC, 1 male OXY+ALC, 1 female OXY+H_2_O). Three rats died during extinction (1 male OXY+ALC, 1 female OXY+H_2_O, 1 male OXY+H_2_O). The SUC+ALC and SUC+H_2_O groups comprised 6 rats of each sex; all completed self-administration. Data from one female SUC+ALC rat was eliminated from analysis as alcohol intake was more than 2 standard deviations from the group mean. Assessment of SSW was not completed for 3 female SUC+H_2_O rats due to experimenter error. BLA vGluT1 signal was undetectable for many oxycodone rats and so only c-fos+ cells were counted in this region.

### Oxycodone, alcohol and sucrose self-administration

There were no Sex x Liquid x Time, Sex x Time, or Sex x Liquid interactions for any oxycodone self-administration variables and thus results are depicted in Fig. 1 without considering sex as a factor. See Fig. S1 for depictions of data accounting for sex. For oxycodone intake, there was a Liquid x Time interaction [F_(11, 341)_=2.301, p<0.01], and the OXY+H_2_O condition displayed greater oxycodone intake than the OXY+ALC condition on Day 10 (Figs. 1b, S1a). The same effect was observed for the total number of infusions earned [F_(11, 341)_=2.406, p<0.01; not shown]. Access to alcohol did not alter active or inactive lever presses during oxycodone self-administration (Figs. 1c, S1b,c), sucrose intake (Figs. 1d, S1d), or active or inactive lever presses during sucrose self-administration (Figs. 1e, S1e,f). Active lever pressing during OXY IVSA [F_(11,341)_=9.660, p<0.0001; Fig. 1c], sucrose intake [F_(11,209)_ =18.130, p<0.0001; Fig. 1d] and active lever presses for sucrose SA [F_(11,209)_=8.435, p<0.0001; Fig. 1e] all increased over the course of the 12 days of SA (main effect of Time). There was a Time x Reinforcer interaction [F_(11, 275)_=1.943, p<0.05] for alcohol intake (g/kg); the SUC+ALC group consumed more alcohol than the OXY+ALC group on Days 10 and 12 (Fig. 1f). For alcohol preference, there was a significant main effect of Reinforcer [F_(1, 25)_=14.196, p<0.001; Fig. 1g], with rats undergoing oxycodone IVSA showing a reduced preference for alcohol relative to sucrose rats.

### Somatic Signs of Withdrawal and Anxiety-like behavior

There was not a Sex x Liquid x Reinforcer interaction on the withdrawal score. There was a Reinforcer x Liquid interaction [F_(1,50)_=8.682, p<0.01], with Oxy+H_2_O rats displaying greater withdrawal signs than all other groups and Oxy+ALC rats displaying withdrawal signs greater than the SUC+H_2_O group (Fig. 2a). SSW were positively correlated with the total amount of oxycodone self-administered in the 12 days prior to assessment [r(35)=0.4991, p<0.01; Fig. 2b], as well as future oxycodone intake on Day 13, 24 hr following SSW assessment [r(33)=0.4219, p<0.05; not shown], but not alcohol intake (not shown).

**Fig. 2.**
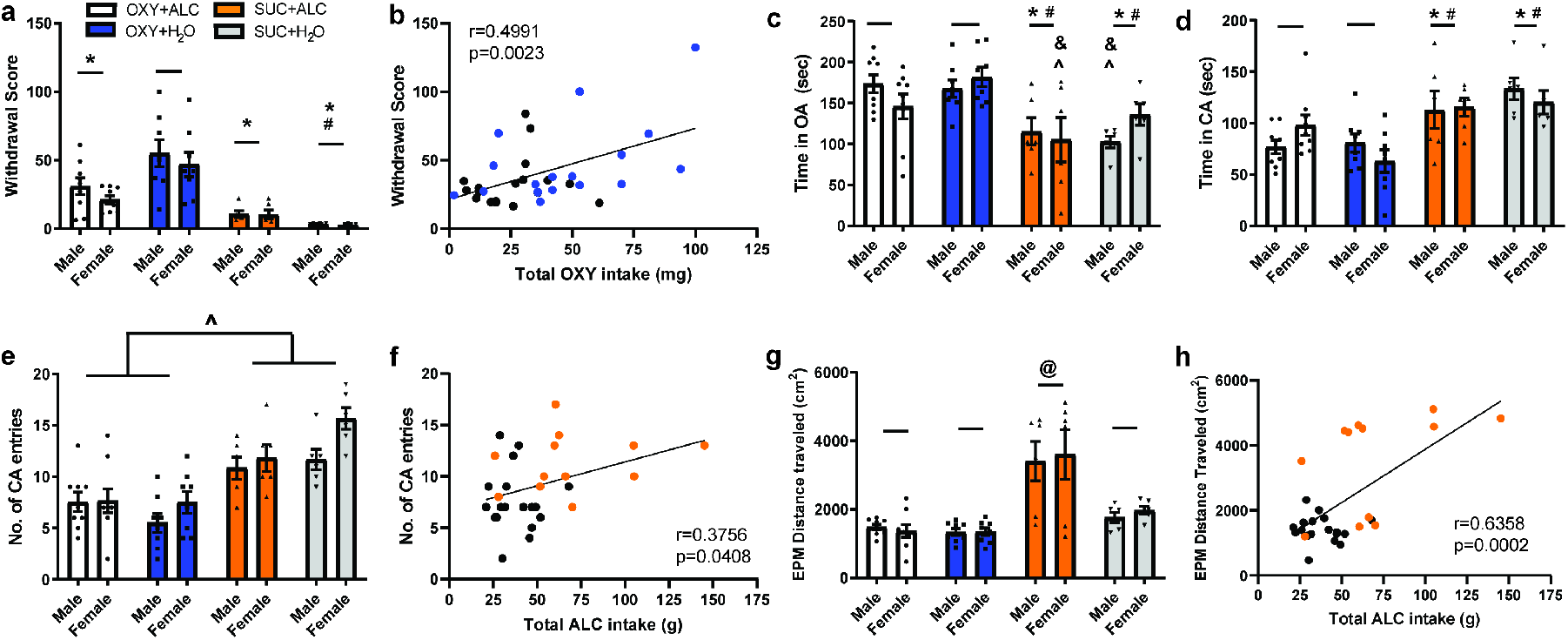
Oxycodone intake increases spontaneous somatic withdrawal signs while alcohol intake is associated with increased anxiety-like behavior. **a.** There were no sex differences in the number of somatic signs of withdrawal exhibited 24 hrs after the last operant session. OXY+H_2_O rats displayed greater withdrawal signs than all other groups and OXY+ALC rats displayed withdrawal signs greater than the SUC+H_2_O group. **b**. The withdrawal score positively correlated with the total amount of oxycodone intake in the 12 days prior to assessment. Blue dots represent OXY+H_2_O rats and black dots are OXY+ALC rats. **c.** Overall, rats that consumed OXY+H_2_O and OXY+ALC spent more time in the open arms (OA) than sucrose-consuming rats, indicating reduced anxiety. **d.** Both sucrose groups spent more time in the CA than the OXY groups. **e.** Sucrose self-administering rats exhibited a greater number of closed arm (CA) entries than OXY self-administering rats. **f.** The number of CA entries was positively correlated with total alcohol intake. Orange dots represent SUC+ALC rats and black dots represent OXY+ALC rats. **g.** The SUC+ALC group displayed greater locomotion than all other groups. **h**. Total alcohol intake was positively correlated with distance traveled in the EPM. * = p<0.05 vs. OXY+H_2_O; # = p<0.05 vs. OXY+ALC; ^ = p<0.05 vs. male OXY+ALC; & = p<0.05 vs. female OXY+H_2_O; @ = p<0.05 vs. all groups.

A Sex x Liquid x Reinforcer interaction was found for time spent in the OA [F_(1,50)_=4.665, p<0.05; Fig 2c]. Overall, OXY+ALC and OXY+H_2_O rats spent more time in the OA than sucrose self-administering rats. Male OXY+ALC rats spent more time in the OA than male SUC+H_2_O and female SUC+ALC rats; female OXY+H_2_O rats spent more time in the OA than male SUC+H_2_O and female SUC+ALC rats. No 3-way interaction was detected for number of OA entries, CA entries, and time spent in the CA. There was a Liquid x Reinforcer interaction for time spent in the CA [F_(1,50)_= 4.620, p<0.05; Fig. 2d], with both SUC groups spending more time in the CA than the OXY groups. There were no 2-way interactions or main effects for number of OA entries (not shown). There was a main effect of Reinforcer for number of CA entries [F_(1,7)_= 42.883, p<0.0001; Fig. 2e], with sucrose self-administering rats exhibiting a greater number of entries. There was a correlation between the amount of alcohol consumed in the 18 days prior to EPM with the number of CA entries [r(30)=0.3756; p<0.05; Fig. 2f]. There was a Liquid x Reinforcer interaction for locomotion in the EPM [F_(1,50)_= 13.724, p<0.001; Fig. 2], with the SUC+ALC group displaying greater locomotion than all other groups (Fig. 2g). Total alcohol intake was positively correlated with distance traveled in the EPM [r(30)=0.6358, p<0.001; Fig. 2h].

### Economic demand for intravenous oxycodone

Demand curve analyses revealed a significant reduction in elasticity of demand (α) for oxycodone in the OXY+ALC group [F_(1, 23)_=10.26, p<0.0001; Fig. 3a]. However, when considering sex, only female OXY+ALC rats displayed less elasticity of demand relative to female OXY+H_2_O rats [F_(1, 18)_=34.81, p<0.0001; Fig. 3b] with no differences in males (Fig. 3c). No group differences were found for Q_0_, essential value, or P_max_.

**Fig. 3.**
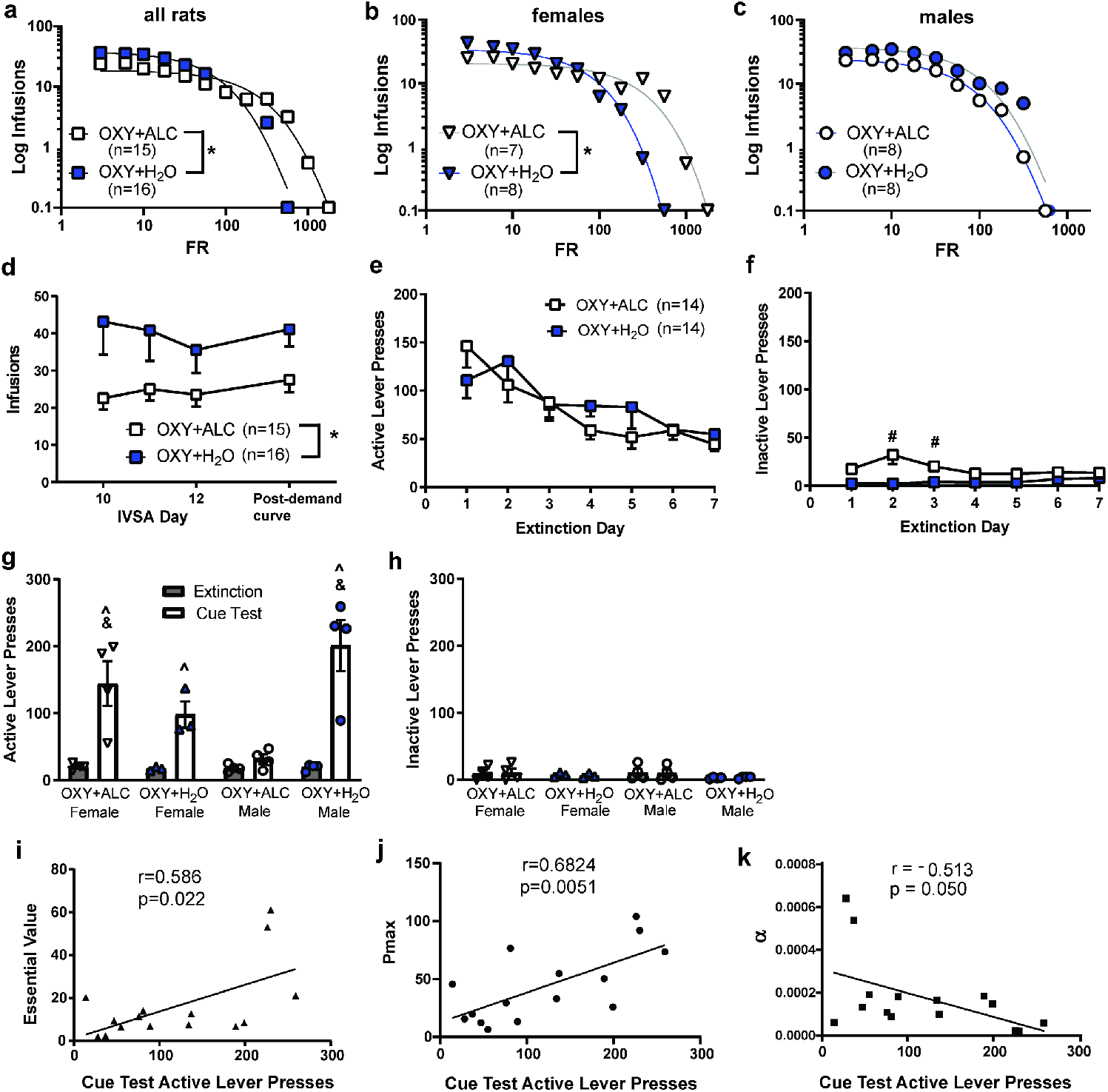
Demand for intravenous oxycodone is increased by co-consumption of alcohol in female but not male rats. **a.** Demand curve analyses comparing all rats found that rats that consumed both oxycodone and alcohol displayed less elastic demand for oxycodone. However, conducting the same analyses separately by sex reveal that this effect is driven by females **(b)** with no group differences evident in male rats **(c). d.** Rats resumed the pre-demand curve level of oxycodone self-administration when placed back onto an FR-3 prior to extinction training and the alcohol-consuming rats attained less oxycodone infusions both before and after the demand curve. **e.** Extinction of oxycodone seeking did not differ by sex; both groups extinguished responding similarly. **f.** Inactive lever pressing was higher for the OXY+ALC group on days 2 and 3 of extinction. **g.** Alcohol co-consumption increases cue-primed reinstatement of oxycodone-seeking in female rats and reduces it in male rats. **h.** Inactive lever pressing during the test was low and did not differ between groups. **i.** Essential Value (α transformed) computed from the oxycodone demand curve analyses was positively correlated with active lever presses during the cue test. **j.** Active lever pressing during the cue test was positively correlated with Pmax derived from oxycodone demand curve analyses. **k**. There was a trend for demand elasticity (α) to be negatively correlated with active lever pressing during the cue test. # = p<0.05 vs. OXY+ H_2_O; ^ = p<0.05 vs. extinction; & = p<0.05 vs. male OXY+ALC. * = effect of Liquid p<0.05.

### Reinstatement of oxycodone-seeking and associated c-fos mRNA expression

Rats readily re-established responding for oxycodone on an FR-3 schedule (Figs. 3d; S1i). To examine whether self-administration behavior was different after the demand curve procedures, we compared the number of infusions attained on the last three days of FR-3 to the amount attained on an FR-3 schedule after the demand curve. There was an effect of Liquid [F_(1, 24)_=5.491, p<0.05] on infusions during this phase, with the alcohol-consuming rats administering less oxycodone infusions both before and after the demand curve. All rats completed at least 7 days of extinction training; during these 7 days, there were no 3- or 2-way interactions on active lever presses and both groups extinguished responding similarly evidenced by a main effect of Time [F_(6, 156)_=10.722, p<0.01; Figs. 3e, S1j]. For inactive lever presses during extinction from oxycodone self-administration, there was no 3-way interaction, but there was a Liquid x Time interaction [F_(6, 156)_=4.817, p<0.05; Figs. 3f, S1k], with overall increased responding in the OXY+ALC condition on Day 2 and 3 of extinction. Only 4 of 7 rats from each condition met extinction criteria by Day 16 and were tested for reinstatement. There was a Sex x Liquid x Time interaction for active lever pressing when comparing the last day of extinction to the reinstatement test day [F_(1,11)=_13.217, p<0.01; Fig. 3g]. Male OXY+H_2_O and Female OXY+ALC rats displayed greater reinstatement (i.e., active lever presses) than male OXY+ALC rats. When comparing lever pressing during the test to those during extinction training, the OXY+H_2_O males and females and the female OXY+ALC groups increased responding during the test, e.g. reinstated responding (p’s<0.01). A paired samples t-test found that the OXY+ALC males only displayed a trend towards an increase in responding from extinction to test (p=0.08). There were no interactions for inactive lever pressing during extinction and test (Fig. 3h). Reinstatement active lever pressing was correlated with several demand curve variables, including reinforcer efficacy, or α transformed [r(15)=0.5859, p=0.0217; Fig. 3i] and with P_max_ [r(15) =0.6824, p<0.01; Fig. 3j]. There was a trend for reinstatement lever pressing to be negatively correlated with α, [r(15)=^-^0.5132, p=0.050; Fig. 3k]. These results indicate a strong relationship between oxycodone-seeking during economic demand procedures and reinstatement tests.

Sites of imaging and representative images are shown in Fig. 4a,b,c. It was first determined which brain regions displayed activation during the cue-test by comparing expression in rats that underwent reinstatement tests with those that did not using 3-way (Sex x Liquid x Test) ANOVAs. When doing this analysis on the number of double-labeled cells (c-fos^+^/vGlut^+^ or c-fos^+^/D1^+^), there was a main effect of Test (i.e. increased c-fos expression in the tested groups) for the IL, NAc, NAs, and dSTR, but not PL (Fig. 4 and Fig. S2). There was an effect of Test on number of c-fos^+^ cells only in the IL, BLA and NAc, indicating that most cells activated by the test are D1-expressing in the NAs and dSTR. There were no 3-way (Sex x Liquid x Test) interactions and only the BLA showed a 2-way interaction (see Supplementary Table 1 for all 3-way ANOVA F stats and p-values).

**Fig. 4.**
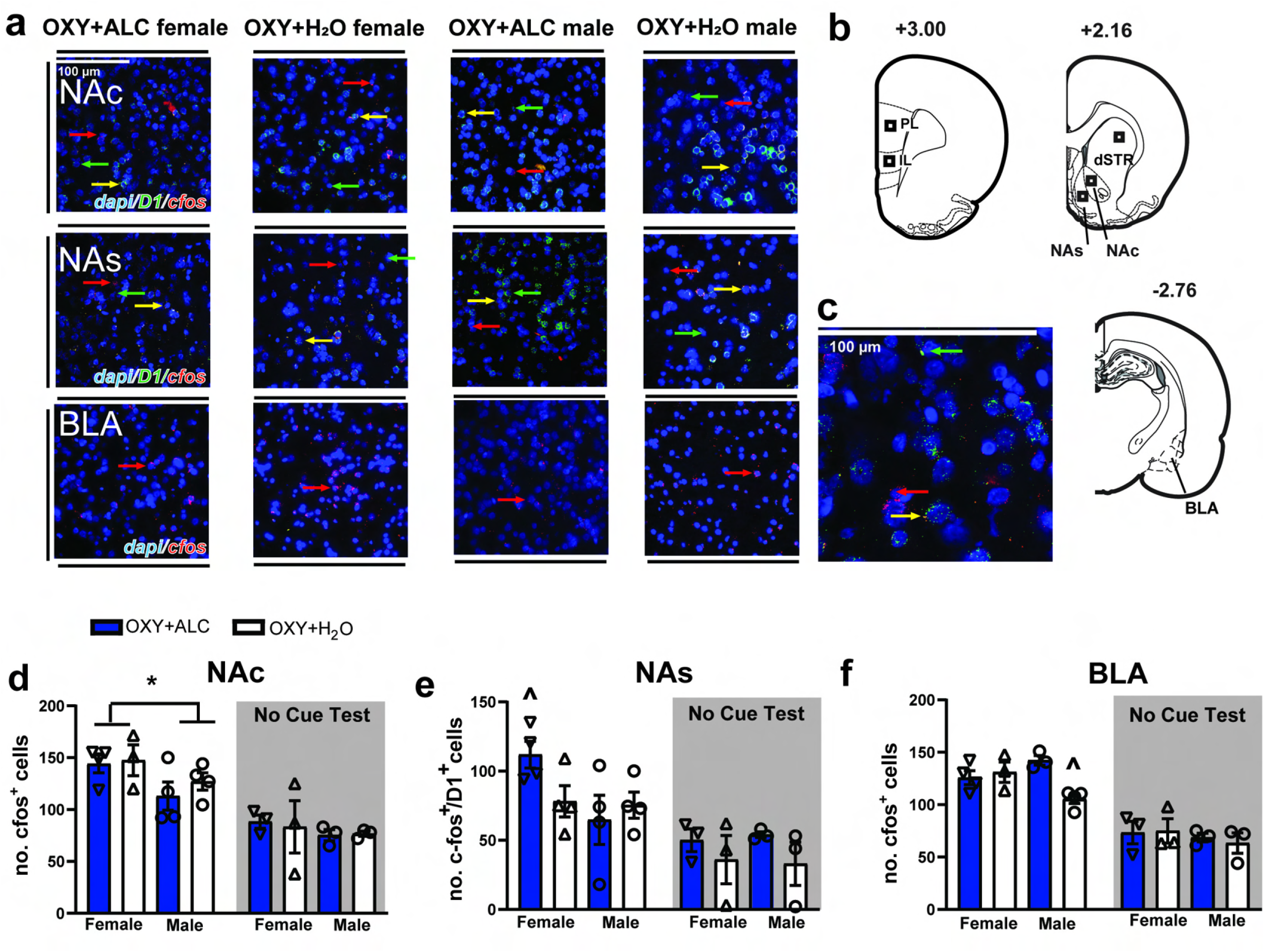
C-fos expression was induced by the cue test and was influenced by alcohol and sex in select brain regions and cell populations. **a.** Representative images from regions and cell populations that showed altered c-fos expression during the test in a manner influenced by a history of alcohol consumption and/or sex. **b**. Sites of analysis in the prelimbic (PL) and infralimbic (IL) cortices, nucleus accumbens (NA) shell (NAs), core (NAc), dorsal striatum (dSTR), and the basolateral nucleus (BLA) of the amygdala. **c.** Representative image illustrating D1/c-fos co-expression in the NAc. **d.** In the NAc, following a cue test, females displayed a greater number of c-fos^+^ cells. There were no group differences in the “no cue test” condition. **e.** In the NAs, following a cue test, females displayed more c-fos^+^/D1^+^ cells than males in the ALC condition. There were no group differences in the “no cue test” condition. **f.** In the BLA, following a cue test, male OXY+ALC rats displayed greater c-fos expression than male OXY+H_2_O rats. There were no group differences in the “no cue test” condition. Red arrows show c-fos mRNA, green arrows indicate D1 mRNA expression and yellow indicates cells where both mRNA are present. *= effect of Sex p<0.05; ^ = p<0.05 vs. OXY+ALC males.

Next, we conducted Sex x Liquid ANOVAs for test and no-test conditions separately, finding no significant effects of Sex or Liquid on c-fos expression in any brain region for the no-test condition, indicating that in the absence of a cue test, c-fos expression does not differ based on sex or a history of consuming alcohol. In the Test condition, there were only effects of Liquid or Sex on expression in the NAc, NAs and BLA. There was a main effect of Sex on the number of c-fos^+^ cells in the NAc [F_(1,11)_=8.816, p<0.05], with females displaying greater number of c-fos^+^ cells (Fig. 4d), an effect that occurred at the trend level for c-fos^+^/D1^+^ cells (p=0.07; Fig. S2a). In the NAs, there was a Sex x Liquid interaction on the number of c-fos^+^/D1^+^ cells [F_(1,11)_=6.117, p<0.05], with females displaying more c-fos^+^/D1^+^ cells than males in the ALC condition (Fig. 4f). There were no such effects on the number of NAs c-fos^+^ cells (Fig. S2b). For the BLA (Fig. 4g), there was a Sex x Liquid interaction on the number of c-fos^+^ cells [F_(1,11)_=10.29, p<0.01], with post-hoc tests finding that male OXY+ALC rats displayed greater activation than male OXY+H_2_O rats. The number of c-fos^+^ cells in the BLA negatively correlated with active lever pressing during the cue test [r(14)= -0.6178, p<0.05].

### Reinstatement of sucrose-seeking and associated c-fos mRNA expression

For sucrose self-administering rats, there was no effect of Sex or any 3-way Sex x Reinforcer x Time interaction for active or inactive lever pressing during extinction or reinstatement (Fig. S2a-d). Active lever pressing decreased during extinction training and was not affected by Liquid [main effect of Time: F_(6, 114)_=99.904, p<0.0001; Fig. 5a]. There was no effect of Liquid or Time on inactive lever presses during extinction (Fig. 5b). To be consistent with methods for the oxycodone self-administering rats, 3-4 rats/sex/condition were tested for sucrose reinstatement. When comparing active lever presses from extinction to the test, there was only a main effect of Time [F_(1,11)_=11.604; p<0.01], indicating that rats reinstated sucrose-seeking without effects of sex or alcohol on this behavior (Fig. 5c). Inactive lever pressing was low and was not influenced by Liquid, but did increase slightly during the test [main effect of Time: F_(1,11)_=6.972; p<0.05; Fig. 5d].

**Fig. 5.**
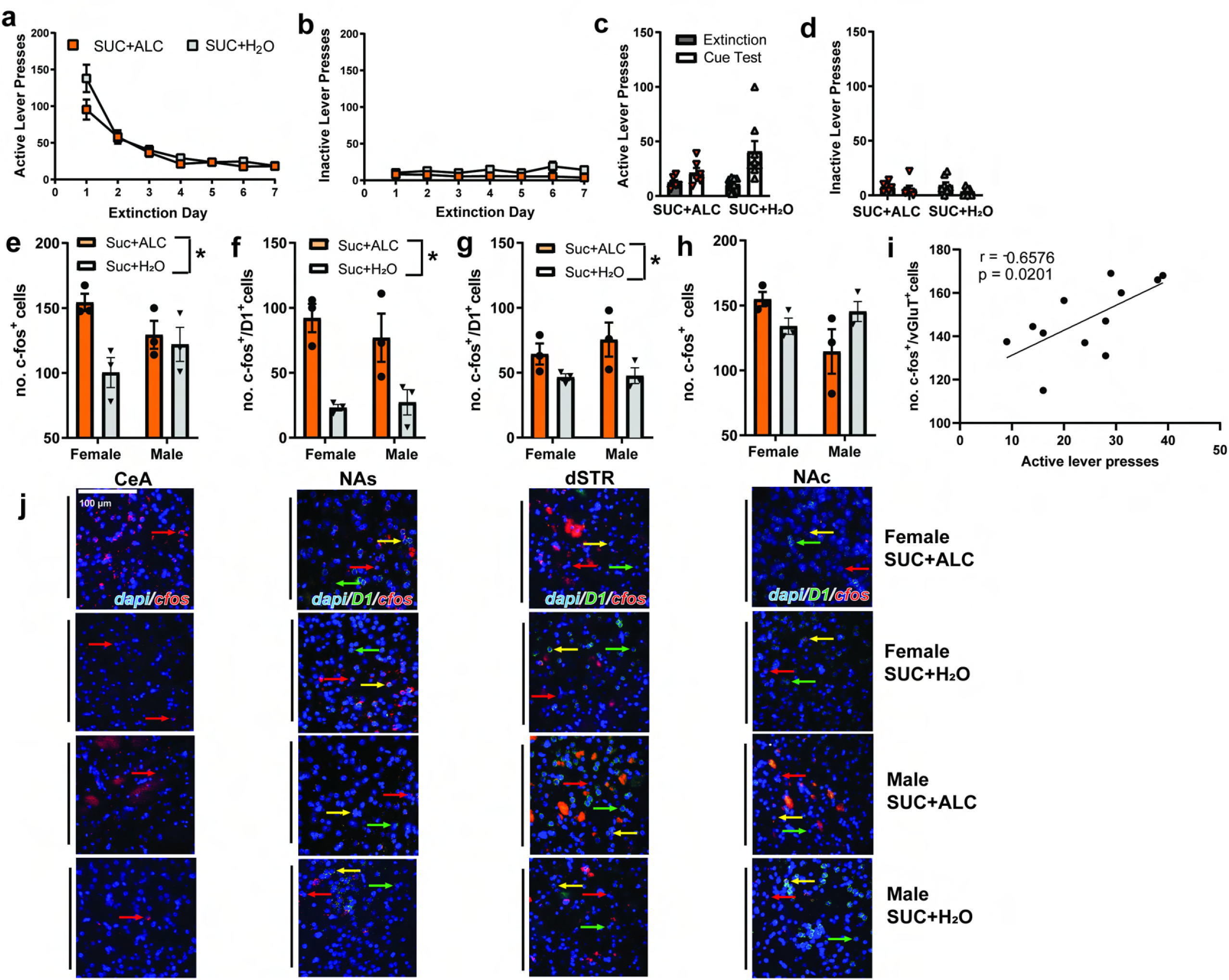
Alcohol intake does not influence lever pressing during extinction and cued reinstatement of sucrose-seeking, but increases c-fos expression following a cue test. **a.** Active lever pressing decreased during extinction training and did not differ between conditions. **b.** Inactive lever pressing did not differ between water and alcohol-consuming groups. **c.** There was a significant effect of Time, but no interactions when comparing lever pressing on the last day of extinction to pressing during the cue test. Males and females are graphed together due to the lack of any effects of Sex on this behavior. **d.** Inactive lever pressing was low and did not differ between groups but did increase slightly during the test. Despite no effects of alcohol on lever pressing during the cue test, alcohol-consuming rats displayed a greater number of c-fos^+^ cells in the CeA **(e)** and more c-fos^+^/D1^+^ cells in the **(f)** NAs and (**g**) dSTR. **h.** There was a Sex x Liquid interaction on c-fos expression in the NAc, with post-hoc tests finding no significant differences between groups. **i**. Only PL c-fos expression correlated with active lever pressing during the test. **j.** Representative images. Red arrows show c-fos mRNA, green arrows indicate D1 mRNA expression and yellow indicates cells where both mRNA are present. * = effect of Liquid p<0.05; & = effect of Sex p<0.05.

There were only 2 sucrose rats/condition that were not tested for reinstatement and thus this tissue was not hybridized. Despite no effects of alcohol on lever pressing during the cue test, there was a main effect of Liquid on the number of c-fos+ cells in the CeA [F_(1, 8)_=8.103, p<0.05; Fig. 5e], and c-fos^+^/D1^+^ co-labeled cells in the NAs [F_(1, 8)_=24.85, p<0.01; Fig. 5f] and dSTR [F_(1, 8)_=7.394, p<0.05; Fig. 5g] with alcohol-consuming rats displaying increased c-fos expression. There was a Sex x Liquid interaction for the number of c-fos^+^ cells in the NAc, with post-hoc tests finding no significant differences between groups [F_(1,8)_=6.327, p<0.05; Fig. 5h]. Only PL c-fos expression correlated with active lever pressing during the test [r(12)=0.6576, p<0.05;Fig. 5i]. Representative images are shown in Fig. 5j.

### Multidimensional Scaling

To gain insight into relationships between the many dependent variables recorded throughout the studies, we conducted MDS analysis. Clusters are depicted in different colors, with the color itself having no significance from panel to panel (Fig. 6). Shorter distances between variables indicate closer correlative relationships and vice versa. Under these analysis conditions, the MDS analysis ignores the direction relationships (i.e. positive vs. negative correlation), showing only where relationships exist. Pearson’s correlations found several correlated variables within each condition/sex (Fig. S4). The optimal number of clusters for each condition was three, but the cluster composition differed by condition (Fig. 6). Relative to female OXY+H_2_O rats (Fig 6a), male OXY+H_2_O rats (Fig. 6b) show several key differences: PL c-fos expression and demand curve variables Q0, EV and log alpha no longer clustering with oxycodone intake, EPM variables no longer clustering with NA c-fos expression, and active lever (AL) presses during the reinstatement test and SSW no longer clustering with AL presses during extinction. Differences in cluster composition between female OXY+H_2_O and OXY+ALC rats are too many to detail (see Fig. 6a and d). Interestingly, female (Fig. 6d) and male OXY+ALC rats (Fig. 6e) display few differences in cluster composition, with only Reinstatement AL presses and SSW belonging in different clusters between the sexes. In both male and female OXY+ALC rats, alcohol intake clustered with NA core and IL c-fos expression, and in male rats only, this cluster also included SSW. For male OXY+H_2_O and OXY+ALC rats, demand curve variables clustered with reinstatement active lever presses, but in OXY+H_2_O rats, this cluster also included c-fos expression in the BLA and NA shell (D1^+^) while in OXY+ALC rats, this cluster included PL c-fos expression. Because sucrose rats have less variables included in the MDS (due to no demand curve analysis), we do not compare SUC to OXY conditions. Alcohol-consuming sucrose rats displayed distinct cluster composition from water consuming rats, with almost no overlap in cluster composition (Fig. 6c, f).

**Fig. 6.**
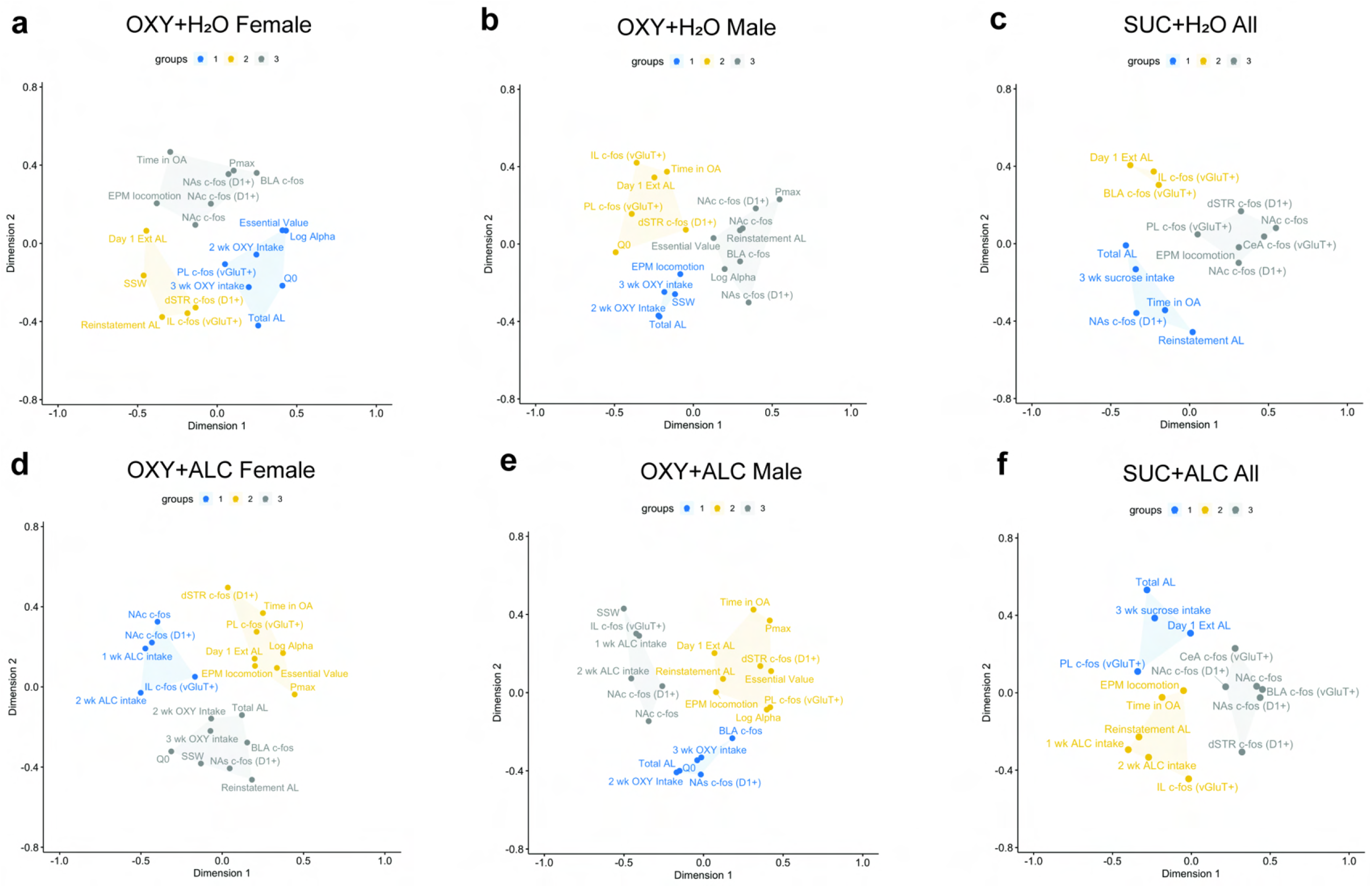
Multidimensional Scaling analysis of key dependent variables. Cluster composition differed by sex for OXY+ H_2_O rats (Panels **a** and **b**) more than it did for OXY+ALC rats (Panels **d** and **e**). Alcohol altered cluster composition in female (Panels **a** and **d**) and male rats (Panels **b** and **e**) that self-administered oxycodone, as well as that of sucrose self-administering rats (Panels **c** and **f**). 2 wk Oxy intake = total oxycodone intake (mg/kg) in the first two weeks of IVSA (all FR-1 and FR-3); 3 wk Oxy intake = total oxycodone intake (mg/kg) in the first three weeks of IVSA (all FR-1, FR-3, FR-6, FR-10, FR-18); 2 wk ALC intake = total alcohol intake (g/kg) in the first two weeks of IVSA; 3 wk ALC intake = total alcohol intake (g/kg) in the first three weeks of IVSA; Total AL = total active lever presses during the first two weeks of IVSA; Time in OA = time spent in the open arms of the EPM; EPM locomotion = total distance traveled in the EPM; Day 1 Ext AL = Day 1 Extinction active lever presses. All other abbreviations defined in main text.

## Discussion

Male and female rats permitted access to both oxycodone and alcohol decreased their consumption of both drugs relative to rats having access to only one substance. However, despite this lower intake, female PSU rats subsequently displayed reduced elasticity of demand for oxycodone that corresponded with increased cue-primed reinstatement of oxycodone-seeking. The opposite pattern was observed for male rats, with reduced reinstatement of oxycodone observed. Spontaneous withdrawal signs were correlated with oxycodone, and not alcohol, intake. Withdrawal from oxycodone was accompanied by reduced anxiety-like behavior in the EPM relative to oxycodone-naïve rats, and alcohol intake was correlated with time spent in the closed arms (e.g., greater anxiety-like behavior). Alcohol access increased the number of CeA neurons activated during cued sucrose-seeking and the number of BLA neurons activated by oxycodone-seeking in males only. Sex-and alcohol-dependent effects on activation of nucleus accumbens D1-expressing neurons were seen following both sucrose and oxycodone reinstatement tests. A data reductionist approach using multidimensional scaling followed by machine-learning based clustering revealed that both sex and alcohol access alter the relationship between key dependent variables. Thus, alcohol alters the motivation to seek oxycodone in a sex-dependent manner and alters the neural circuitry engaged by cue-primed reinstatement of sucrose and oxycodone-seeking.

### Drug intake

Contrary to our hypotheses, the PSU condition reduced intake of oxycodone and alcohol relative to monosubstance conditions. While this could be due to cross-sensitization, only limited evidence supports this possibility; repeated morphine treatment results in cross-sensitization to the locomotor effects of alcohol, however the converse is not true [41]. The effects of non-contingent morphine on alcohol intake are biphasic, with low doses increasing voluntary alcohol intake and high doses decreasing intake; the decrease was blocked by co-administration of naloxone, indicating that signaling through the mu opioid receptor underlies this effect [42]. It is possible that oxycodone-induced desensitization of the mu opioid receptor reduced motivation to consume alcohol, as systemic antagonism decreases alcohol consumption [43]. While such neurochemical adaptations may underlie the effects of oxycodone on alcohol intake, it’s less clear why alcohol would alter oxycodone intake on the following day. Alcohol access had no effect on sucrose intake, in agreement with prior work [44], and ruling out generalized effects on motivation to seek rewards. It is possible that there are pharmacokinetic interactions between the drugs. The half-life of intravenous oxycodone is approximately 62-83 minutes in the rat[45], and thus was likely still present at the time rats were presented with alcohol. It is possible that oxycodone altered the pharmacokinetics of alcohol to decrease intake. Thus, while there is evidence for alcohol–opioid interactions in the literature, particularly at the mu opioid receptor, the underlying pharmacokinetic and pharmacodynamic mechanisms behind the ability of alcohol to alter the motivation to seek oxycodone is currently unclear and warrants further study.

### Somatic signs of withdrawal and anxiety-like behavior

While withdrawal from both alcohol and oxycodone has the potential to result in SSW, there was not an additive effect of the two drugs on spontaneous withdrawal signs. Alcohol intake in sucrose rats did not increase SSW relative to the drug-naïve condition. Noncontingent alcohol exposure and high levels of intake in mice (e.g. 10 g/kg/day) reliably produce SSW; however, such withdrawal signs are modest in models where blood alcohol levels (BALs) range from 150-200 mg% [18]. While we did not assess BAL here, in our previous work employing 6 hr home cage access to 20% alcohol, achieving a range of intake (2-4 g/kg) similar to that observed here, BALs were in the range of 50-80 mg% over the first 2 hours of drinking [23]. Thus, the present model likely does not engender high enough intake to produce SSW. We are the first to report that that SSW are positively correlated with oxycodone intake on the days prior to and after the assessment of SSW, in agreement with results found following heroin IVSA in male and female rats [14]. Following oxycodone IVSA, other measures of withdrawal-related aggression and hyperalgesia have been reported [46].

At the group level, alcohol intake increased locomotion in the EPM but did not alter arm entries. However, the number of closed arms entries was positively correlated with alcohol intake. This may have been a result of the increased locomotion, and not increased anxiety. Future work should explore additional assay to confirm that alcohol withdrawal is anxiogenic and promotes hyperlocomotion, in agreement with previous work [47]. Interestingly, the groups that consumed oxycodone displayed the least anxiety-like behavior in the EPM, with no additive effects of alcohol intake, potentially identifying anxiolysis as a factor underlying the motivation to consume both drugs.

### The effects of alcohol on oxycodone demand, reinstatement of sucrose-and oxycodone-seeking, and associated brain activation

While alcohol co-use decreased oxycodone intake during training in both sexes, in female rats, elasticity of demand for oxycodone was decreased in oxycodone+alcohol rats. We employed a model in which FR requirements for IV drug increased, as opposed to other models which use drug concentration “thresholding” procedures. Here, the motivation to seek oxycodone was assessed when rats were largely undrugged (e.g., needed to emit lever presses to receive the first infusion) and thus acute pharmacological tolerance or sensitization does not explain these results. Demand curve variables (Pmax, essential value) were correlated with active lever presses during the undrugged cue-primed reinstatement test, in agreement with similar work with cocaine and methamphetamine [48,49]. Together, these data indicate that in females, opioid-alcohol co-consumption increases the motivation to seek oxycodone and the reinstatement of oxycodone-seeking. Alcohol did not alter demand elasticity in males, but cue-primed reinstatement of oxycodone-seeking did not occur when male rats had a history of consuming alcohol and oxycodone. However, it should be noted that there was an almost 2-fold increase in lever presses from extinction to test which not attain statistical significance. Alcohol co-consumption did not alter the reinstatement of sucrose-seeking. Home cage alcohol consumption has also been demonstrated to increase Q_0_ (preferred consumption when the price is zero) for intravenous cocaine, without affecting demand elasticity [50].

Despite the relatively small sample size, several brain regions showed increased neuronal activation (e.g., number of c-fos^+^ cells) upon oxycodone reinstatement testing relative to untested rats: the NA core and shell, dSTR, BLA and IL cortex. In the NA shell and dSTR, these increases were restricted to D1^+^ neurons, consistent with a role for these neuronal populations in driving drug-seeking [37]. The PL did not show a test-induced increase, but instead showed overall high expression in both tested and untested conditions, indicating the potential for lasting effects of oxycodone and alcohol intake on PL activity. Sex and alcohol-dependent changes in c-fos expression following oxycodone reinstatement were observed only in three regions, the NA core, NA shell, and BLA. In the NA shell, females in the OXY+ALC group displayed the greatest amount of reinstatement-induced activity, specifically in D1^+^ cells. Cocaine reinstatement-induced c-fos expression is greater in females than males in the NA core and shell, but only when the cocaine-cue pairings occurred during the estrus phase of the estrous cycle [51]. Here, reinforcer-cue pairings occurred throughout the estrous cycle and possibly contributed to greater cue-induced c-fos mRNA expression in females. Male OXY+ALC rats displayed reduced test-induced c-fos expression in the NA core, and decreased reinstatement relative to other groups, indicating that this region may be impacted by PSU and be involved in mediating relapse. Active lever pressing during the cue test was negatively correlated with the number of c-fos^+^ cells in the BLA. A role for this brain region in the inhibition of reinstatement for any drug has not yet been reported in the literature. Taken together, unlike the profound effects of alcohol on cocaine reinstatement-induced c-fos expression observed in these regions in a similar sequential model of cocaine-alcohol PSU, only modest effects were observed here. Thus, it may be the reinstatement of oxycodone seeking relies on similar circuitries in the presence and absence of alcohol consumption.

A history of alcohol consumption increased the number of CeA neurons activated during cued sucrose-seeking, and the number of ventral and dorsal striatum D1-expressing neurons engaged by sucrose reinstatement, without an increase in sucrose-seeking. Reinstatement-induced c-fos expression may be enhanced by increased baseline synaptic activity following a history of alcohol intake for 3 weeks or greater.

Increases in glutamate transmission and/or decreases GABA interneuron activity in the BLA and dorsal and ventral striatum have been identified following alcohol exposure [52–55]. The small number of sucrose rats in the untested condition did not permit assessment of the long-term effects of alcohol alone on c-fos expression. However, following 2 months of IAA, male and female Sprague-Dawley rats displayed no changes in NA core, shell, CeA and BLA c-fos expression at 28 days withdrawal, relative to alcohol-naïve rats [56]. Thus, it is likely that the interaction between alcohol and the cue test increased c-fos expression in these regions.

### Sex differences in oxycodone self-administration and seeking

We observed no sex differences in oxycodone intake or self-administration behavior in both oxycodone-only and PSU groups. This is consistent with work done in Sprague Dawley rats as well as in other strains of rats (F344/N, ACI/N and WKY/N)[45,57]. However, there have been reports of female rats (Wistar) self-administering a greater number of infusions than males [46]. Despite no differences in oxycodone intake, Sprague Dawley females have been found to have higher cued seeking after 14 days of home cage abstinence [57], while no sex differences in this measure were found after 4 weeks of abstinence [45]. Here, in the oxycodone-only rats, there was no statistical difference in active lever presses during the cue-primed reinstatement test. A shorter drug-free period was used here, as was instrumental extinction, which could explain differences in results. In sum, sex differences in oxycodone intake and seeking seem to be dependent on the strain of rat and addiction model used.

### Multidimensional Scaling Analysis

MDS analysis revealed that sex had less of an effect on the cluster composition than did access to alcohol. There are a greater number of differences in cluster composition between female OXY+H2O and female OXY+ALC rats than similarities. Interestingly, male and female OXY+ALC rats display few differences in cluster composition, despite markedly different effects of alcohol on economic demand for oxycodone and the reinstatement of oxycodone-seeking in the sexes. Similarly, while inferential statistics reveal no differences between OXY+H_2_O and OXY+ALC male rats in terms of demand parameters and c-fos expression in many brain regions, MDS-based clustering finds distinct clusters in these conditions, with the BLA, NA core and shell c-fos expression clustering with active lever presses during reinstatement in the OXY+H_2_O males, and the dSTR and PL c-fos expression clustering with this behavior in the OXY+ALC males. While alcohol intake could underlie the differences in cluster composition, alcohol also influenced other dependent variables (e.g. oxycodone intake), which may in turn alter the relationships. These findings highlight the importance of utilizing multiple analysis approaches to decipher complex relationships, including those involving multiple drugs and sexes.

## Conclusions

Alcohol consumption reduces oxycodone intake during self-administration in both sexes, while altering the motivation to seek oxycodone in a sex-dependent manner. Inferential statistics found that alcohol-oxycodone PSU has modest effects on the neural circuitry engaged by cue-primed reinstatement of oxycodone-seeking.. The MDS analysis found that alcohol intake altered the relationship between oxycodone-seeking variables (from intake to economic demand and reinstatement), c-fos expression, anxiety-like behavior and withdrawal signs. Sex also had an impact on such relationships, but more so in the oxycodone-only condition and less in the PSU condition. These results build upon our prior work finding that sequential cocaine-alcohol self-administration alters the role of accumbens glutamate release in cocaine reinstatement as well as reinstatement-induced c-fos protein expression throughout the reward circuitry [23,58]. Future work examining pharmacological interactions between alcohol and oxycodone are necessary to interpret the ability of these drugs to influence the intake of one another. These data, together with the clinical significance of PSU, strongly indicate that more research is needed to characterize the behavioral and neurobiological effects of PSU in order to develop targeted therapies for reducing drug use.

## Author contributions

LAK, MS, and CSW designed the studies and analyzed the behavioral data. CSW, GR and LW conducted the behavioral studies. LW conducted the histology and in situ hybridization. HLB and SD imaged and conducted analysis of mRNA expression. NPM assisted in data analysis. All authors contributed to the writing of the manuscript.

## Data Availability Statement

Data will be made available upon request following publication.

