## Supplementary Materials for "Voluntary alcohol intake alters the motivation to seek intravenous oxycodone and neuronal activation during the reinstatement of oxycodone and sucrose seeking"

### Supplementary Data

**Supplementary Table 1.** Results of 3-way Sex x Liquid x Test ANOVAs for c-fos expression in Oxy rats.

| **Dependent Variable** | **Main effect of Test** | | **Sex x Liquid x Test** | | **Liquid x Test** | | **Sex x Test** | |
|---|---|---|---|---|---|---|---|---|
| | F stat | p-value | F stat | p-value | F stat | p-value | F stat | p-value |
| IL # c-fos$^+$/vGlut1$^+$ cells | 11.39 | **<0.05*** | 0.52 | n.s | 0.04 | n.s | 1.48 | n.s |
| PL # c-fos+/vGlut1+ cells | 3.04 | n.s | 0.78 | n.s | 0.19 | n.s | 0.55 | n.s |
| NAc # c-fos$^+$/D1$^+$ cells | 28.49 | **<0.001*** | 0.15 | n.s | 0.04 | n.s | 1.04 | n.s |
| NAs # c-fos$^+$/D1$^+$ cells | 17.56 | **<0.001*** | 1.75 | n.s | 0.12 | n.s | 0.18 | n.s |
| dSTR # c-fos$^+$/D1$^+$ cells | 11.27 | **<0.01*** | 3.07 | n.s | 0.55 | n.s | 0.20 | n.s |
| IL # c-fos$^+$ cells | 6.03 | **<0.05*** | 1.48 | n.s | 0.02 | n.s | 0.87 | n.s |
| PL # c-fos+ cells | 2.03 | n.s | 0.49 | n.s | 0.01 | n.s | 0.45 | n.s |
| BLA # c-fos$^+$ cells | 99.55 | **<0.05*** | 2.46 | n.s | 4.80 | **<0.05*** | 0.11 | n.s |
| NAc # c-fos$^+$ cells | 35.58 | **<0.05*** | 0.02 | n.s | 0.37 | n.s | 0.83 | n.s |
| NAs # c-fos$^+$ cells | 3.31 | n.s | 2.16 | n.s | 0.03 | n.s | 0.01 | n.s |
| dSTR # c-fos$^+$ cells | 2.63 | n.s | 1.71 | n.s | 0.002 | n.s | 0.01 | n.s |

### Supplementary Figures

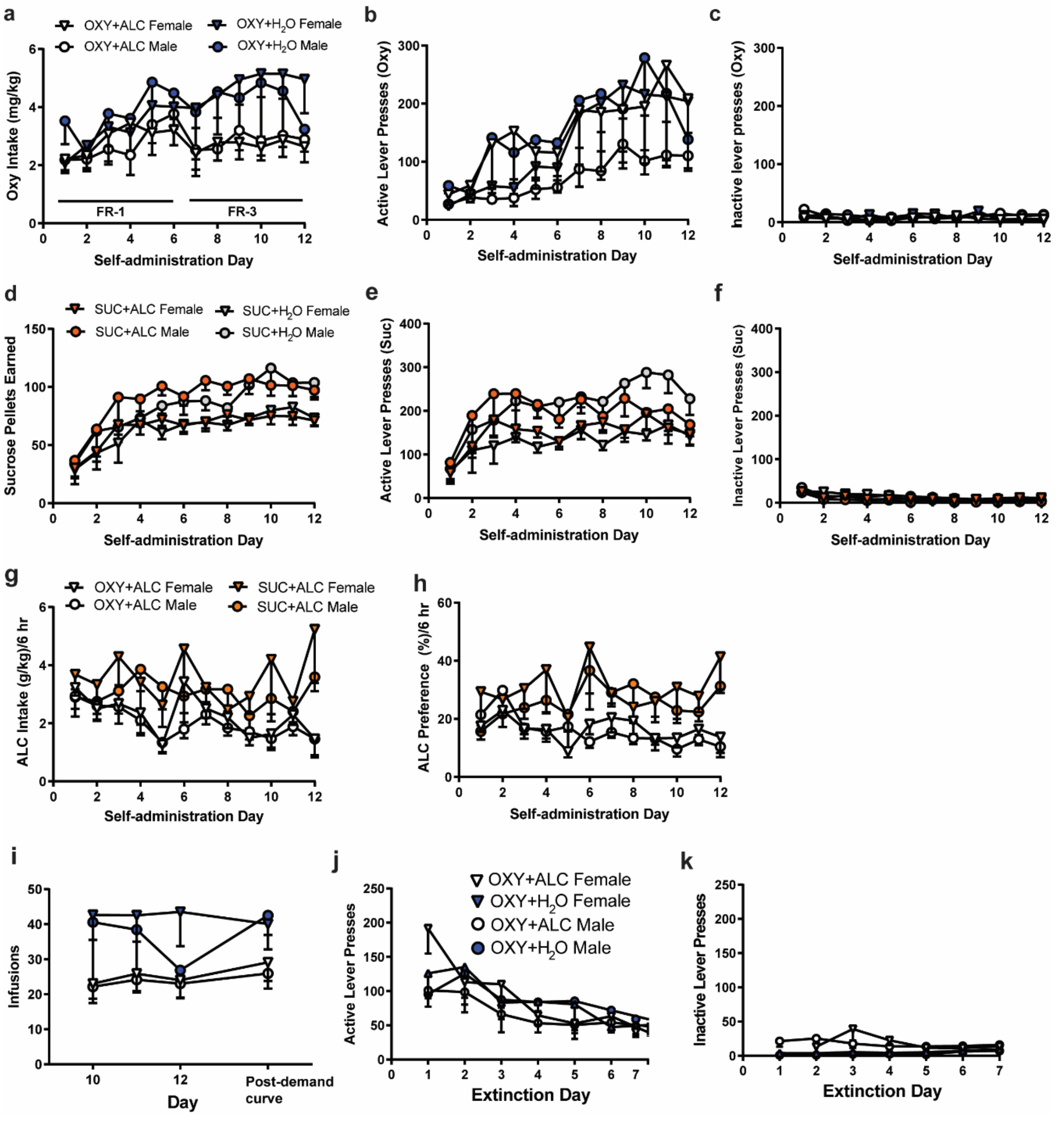

**Supplementary Fig. 1.** There were no effects of sex on oxycodone intake (**a**), active lever presses (**b**), or inactive lever presses (**c**). There were no effects of sex on sucrose intake (**d**), active lever presses (**e**), or inactive lever presses (**f**). There were no sex differences in alcohol intake (**g**) or preference (**h**). **i.** There were no effects of sex on oxycodone intake prior to and after the demand curve. There were no effects of sex on active (**j**) or inactive lever presses (**k**) during extinction training.

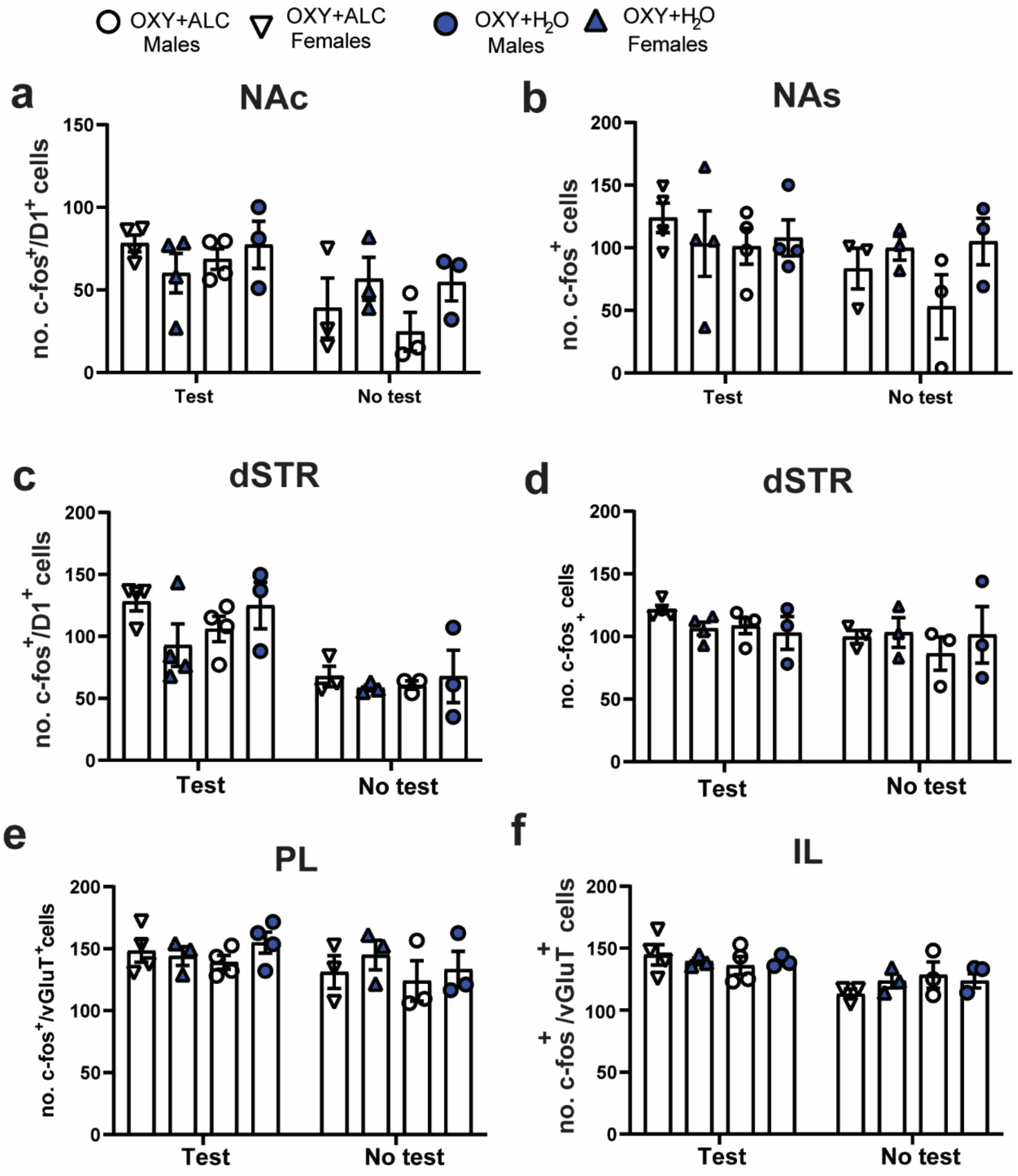

**Supplementary Fig. 2**. **a.** There was a main effect of Test on c-fos expression in D1$^+$ cells of the NAc, but no effects of Liquid and a trend for an effect of Sex (p=0.07) on expression. **b.** There was no effect of Test on c-fos$^+$ cells in the NAs. **c**. There was a main effect of Test for c-fos expression in D1$^+$ cells of the dSTR, expression did not differ by Sex or Liquid. **d**. There was no effect of Test on c-fos$^+$ cells in the dSTR. **e**. There was no effect of Test on c-fos expression in vGlut1$^+$ cells of the PL. **f.** There was a main effect of Test for c-fos expression in vGlut1$^+$ cells of the IL, but no effects of Liquid or Sex on expression.

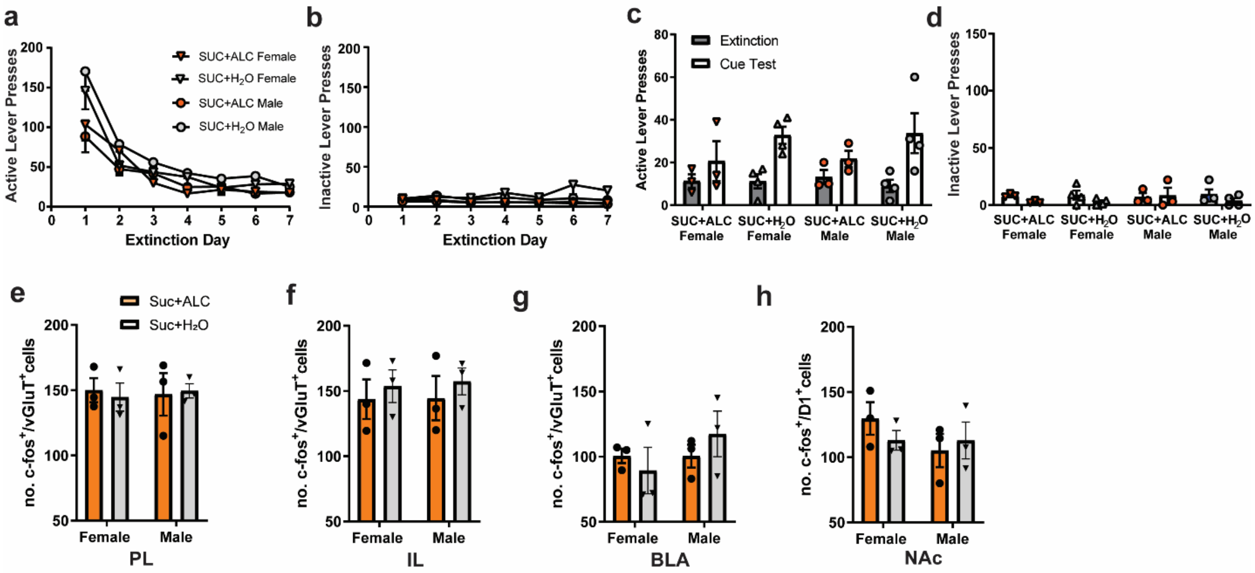

**Supplementary Fig. 3.** There were no effects of sex on active lever presses (**a**), or inactive lever presses (**b**) during extinction training following sucrose self-administration. There were no effects of sex on active lever presses (**c**), or inactive lever presses (**d**) when comparing the last day of extinction training to the cue test. Alcohol did not alter c-fos expression following the sucrose cue-test in male or female rats in glutamate neurons of the PL (**e**), IL **(f)**, BLA (**g**), or in D1-expressing neurons in the NAc (**h**).

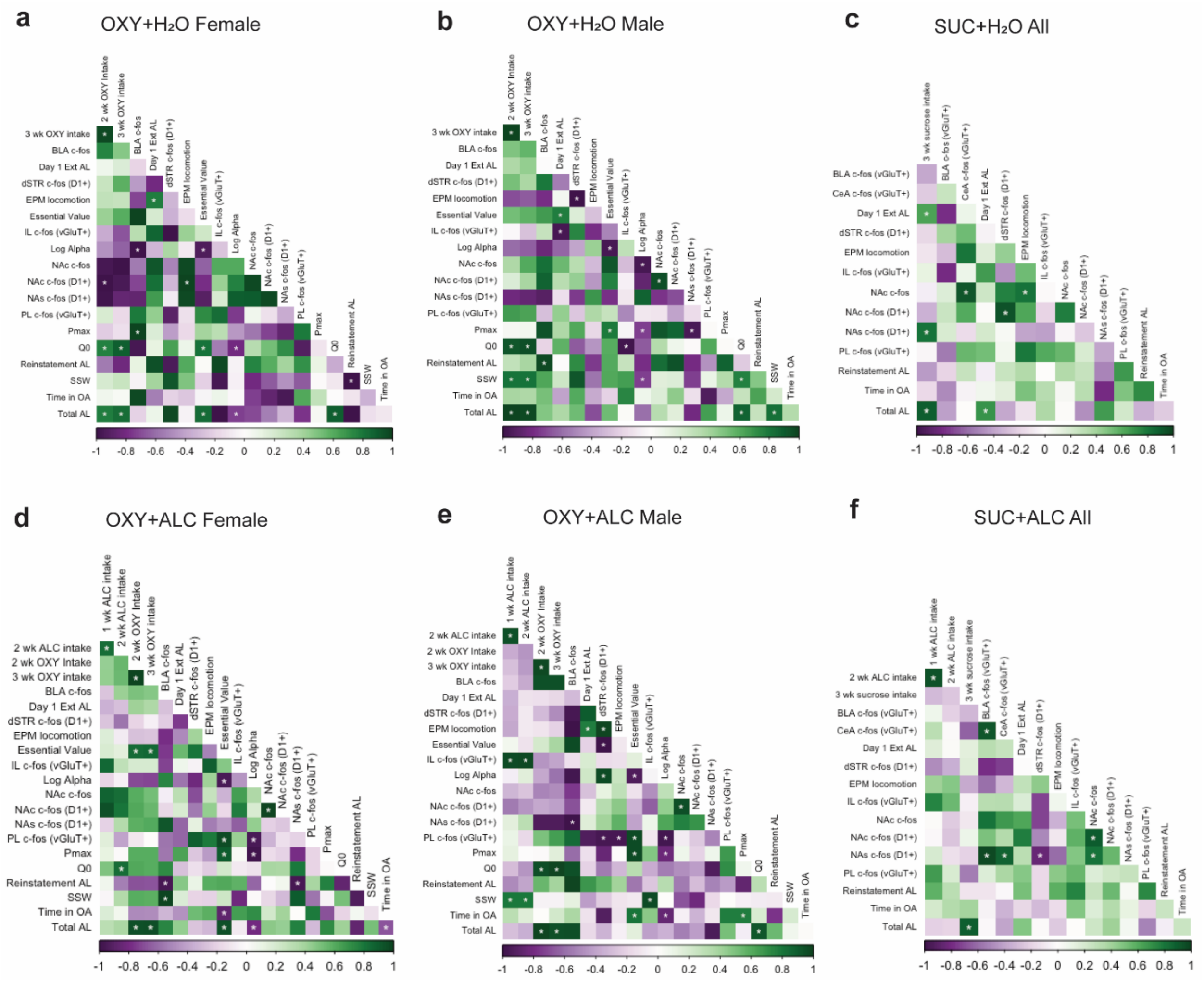

**Supplementary Fig. 4.** Pearson's correlations. * = $p<0.05$
